# Single Capture Quantitative Oblique Back-Illumination Microscopy

**DOI:** 10.1101/2025.07.29.667497

**Authors:** Paloma Casteleiro Costa, Srinidhi Bharadwaj, Zhenmin Li, Nischita Kaza, Mercedes Lopez-Esteva, Anthony Lien, Bilal Haider, Francisco E. Robles

**Author notes:** Contributing authors.

## Abstract

Quantitative oblique back-illumination microscopy (qOBM) has emerged as a powerful technique for label-free, 3D quantitative phase imaging of arbitrarily thick biological specimens. However, in its initial embodiment, qOBM requires multiple captures for phase recovery, which reduces imaging speed and increases system complexity. In this work, we present a novel advancement in qOBM: single-capture qOBM (SCqOBM) which utilizes a deep learning model to accurately reconstruct phase information from a single oblique back-illumination capture. We demonstrate that SCqOBM achieves remarkable phase imaging accuracy, closely matching the results of traditional four-capture qOBM in diverse biological samples. We first highlight the unique potential of SCqOBM for non-invasive, in-vivo imaging applications by visualizing blood flow in mouse brain and human arm. Additionally, we demonstrate single-slice (en-face) quantitative phase imaging at 2 kHz and volumetric refractive index tomography at speeds up to 10 volumes per second. SCqOBM offers transformative advantages in speed, simplicity, and system accessibility, making it highly suitable for dynamic and real-time imaging applications. Its ability to produce high-resolution, quantitative phase and refractive index images with minimal hardware complexity opens new frontiers in biomedical research and clinical diagnostics, including non-invasive hematological assessments and in-vivo tissue imaging.

## 1 Introduction

There has been a growing need for fast, label-free imaging technologies that can provide clear cellular and subcellular contrast for a wide range of medical and biomedical applications. To date, a number of very promising technologies exist for this purpose, but trade-offs vary widely. Optical coherence tomography (OCT), for example, provides extremely fast (video rate) volumetric imaging [1, 2], but its spatial resolution and contrast to cellular and subcellular structures is low. Label-free nonlinear microscopy (e.g., Coherent Raman Scattering [3], Pump Probe microscopy [4], Simultaneous Label-free Autofluorescence Multi-harmonic microscopy [5]) provides superior resolution and (sub)cellular detail, but such methods are much slower due to the need for laser point scanning. Plus, technologies that require ultra-fast pulses are complex, expensive and can cause tissue damage; consequently, they have not been widely adopted for in vivo clinical imaging applications. Finally, label-free autofluorescence light sheet microscopy is extremely fast [6], but has limited contrast, and the penetration depth is low (∼ 50 *µm*). To overcome these trade-offs, we introduce single-capture qOBM (SCqOBM). This technology achieves quantitative phase and refractive index information with high-resolution, achieves an imaging depth similar to nonlinear microscopy, and provides single-shot 2D multiplexing, like light sheet microscopy. We demonstrate that SCqOBM achieves clear cellular and subcellular contrast at imaging rates of up to 2 kHz for a single slice in 3D space or at near video-rates (10 Hz) for volumetric imaging, thus enabling new opportunities for label-free optical imaging across a wide range of applications.

qOBM is a significant advancement of quantitative phase imaging (QPI), so to understand this technology and its unique capabilities, first consider QPI which has become an essential tool for biology, biomedicine and bioengineering. QPI enables observation of live, unlabeled specimens to study the composition of complex cellular and subcellular structures at the nanometer scale, as well as dynamic phenomena such as cell proliferation and cell-mass transport with unparalleled sensitivity[7–9]. Traditional QPI methods, however, rely on light being transmitted through samples to measure phase delays, limiting their application to thin specimens (e.g., thin monolayer cell cultures). Even with the more recent advent of 3D QPI (also known as optical diffraction tomography, ODT, or refractive index tomography) [10, 11], which can resolve multiple cell layers, the technology remains a transmission-based technique. Consequently, traditional QPI and ODT methods cannot be used to study large specimens (typically thicker than 1-2 mean-free-scattering path lengths).

qOBM was introduced to overcome this significant limitation of QPI [12– 15]. We have recently demonstrated that qOBM enables high-resolution, label-free quantitative phase imaging and refractive index tomography with the same rich quantitative information as traditional QPI and ODT but now also in arbitrarily thick samples–which were previously inaccessible–by using epi-illumination instead of transmission-based illumination, a non-trivial advancement.

Reconstruction for qOBM images follows the principles of oblique illumination with partially coherent light to recover phase and refractive index information in 3D [15–17]. Specifically, in qOBM four raw image captures are acquired to first produce two pairs of orthogonal differential phase contrast (DPC) images (Figure 1A-B).The DPC images are subsequently used to reconstruct a quantitative phase image of a slice in 3D space via deconvolution (Figure 1B). Alternatively, a stack of DPC images can be acquired and deconvolved to directly obtain the 3D refractive index distribution [15]. This approach is relatively fast and simple to implement and has been used to study 3D cell culture dynamics [13, 18], organoids [19], cord blood unit viability [14], brain tumor pathology [20, 21], and root microbe dynamics [22], among other applications. Despite its demonstrated utility, qOBM has one main limitation: the need for four captures to reconstruct the quantitative phase information, which (1) reduces the maximum imaging rate to one-fourth the camera’s frame rate, (2) subjects the system to motion artifacts when analyzing fast dynamics, and (3) increases system complexity.

**Fig. 1.**
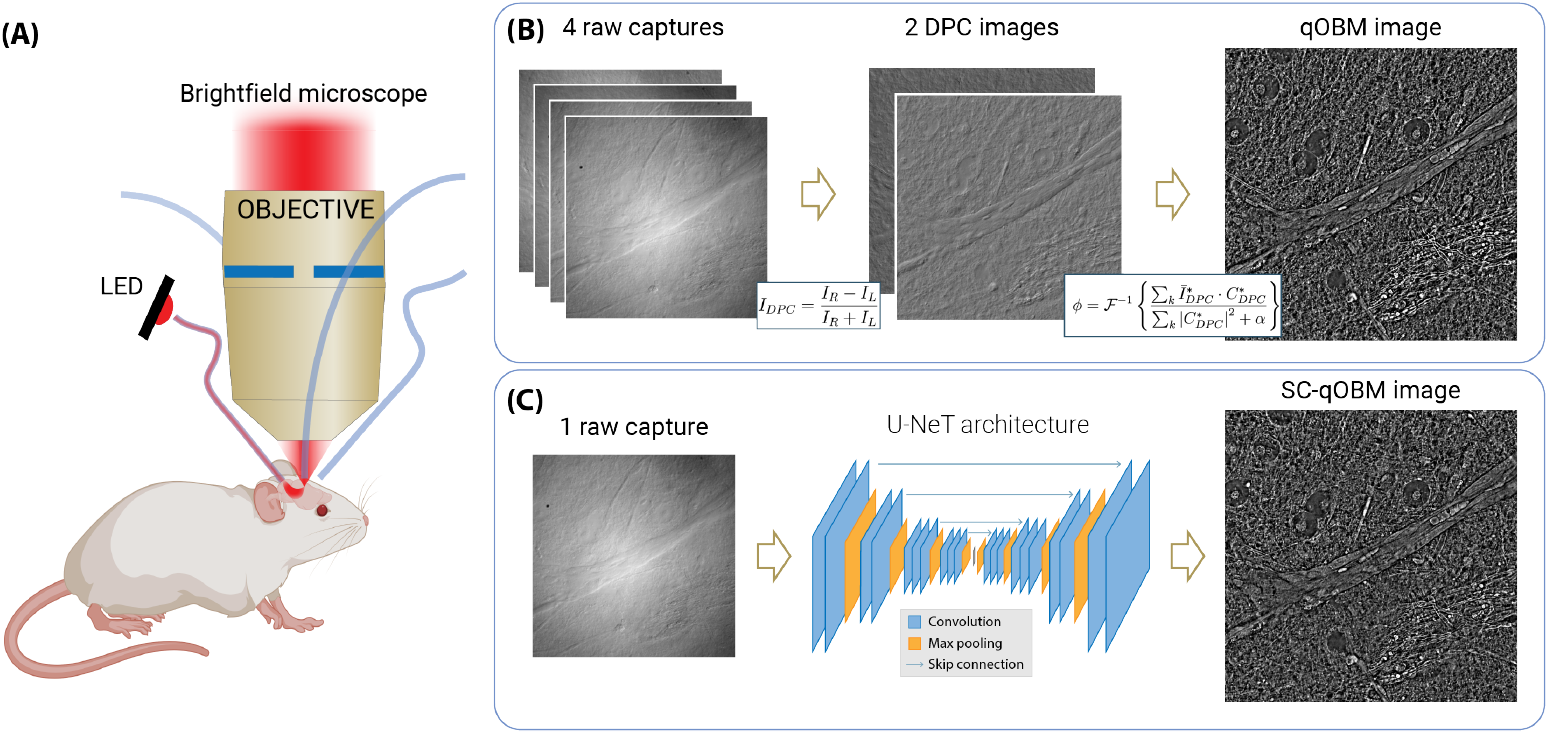
qOBM and SCqOBM system and phase retrieval process. (A) System schematic. (B) In qOBM four captures are acquired by illuminating the sample from four oblique angles. These captures are then used to obtain two differential phase contrast images which are then used to reconstruct a qOBM image through a Tikhonov deconvolution. (C) in SCqOBM, one OBM capture is fed through a U-Net model trained to directly retrieve the quantitative phase.

To improve the imaging speed of qOBM, this work introduces SCqOBM which leverages recent advances in deep learning (DL), a powerful technique within artificial intelligence (AI) that has garnered considerable attention across various disciplines, including microscopy and medical imaging. DL has shown remarkable success in performing complex tasks such as data classification [20, 23], data segmentation [24], particle tracking [25], image enhancement [26, 27], automated real-time focusing [28, 29], and virtual staining of label-free data [30–32]. Here we demonstrate that the DL-enabled, SCqOBM achieves high-fidelity reconstruction of quantitative phase tomograms from a single oblique back illumination capture. We also develop and compare two-capture (TC)qOBM, which uses two orthogonal oblique back illumination captures as input. We train and validate the approach using static samples, where ground truth is known *apriori*. Then, we highlight the unique capabilities of the approach by imaging blood flow in-vivo in a mouse brain window chamber model and in human skin. We also demonstrate qOBM imaging in a single plane (en-face slice) at rates of 2 kHz and qOBM volumetric refractive index tomography rates of 10 volumes per second in vivo.

This novel qOBM approach effectively achieves similar imaging speeds as light sheet microscopy (both achieve 2D spatial multiplexing with a single capture), but with similar resolution and imaging depth as label-free non-linear microscopy and with the unique quantitative contrast and subcellular detail of QPI. The technology also uses simple and low-cost instrumentation (bright field microscope with epi-illumination using one LED). Together, SCqOBM (and TCqOBM) significantly broadens the utility of fast, label-free optical imaging which could have significant implications for many clinical, pre-clinical and high throughput biomedical applications.

## 2 Results

### 2.1 Fast qOBM of static (*ex-vivo*) samples

#### 2.1.1 Umbilical cord blood inside collection bags

The SCqOBM and TCqOBM models were first trained and tested using a dataset comprising images of umbilical cord blood inside collection bags, as decried in Ref. [14]. This specific dataset provides predictable, non-dynamic structures (red blood cells (RBCs) and white blood cells (WBCs)) to first test and evaluate the feasibility and performance of SC- and TCqOBM with idealized, yet real, samples. The data were acquired with a dual-wavelength configuration, where one of the LED pairs had a central wavelength of 530 nm (for the first DPC image), and the other pair had a central wavelength of 627 nm (for the orthogonal DPC image) (see Methods Section). (Note that two colors were used here to differentiate between WBC and RBC using structure and hemoglobin absorption.) The TCqOBM model’s input consisted of two raw captures, one from each set of orthogonal DPC images (one at 530 nm and the other at 627 nm). For SCqOBM, the model’s input comprised only one 627 nm acquisition. Model architecture and training parameters are described in the Methods Section.

Figure 2 illustrates a set of images of cord blood inside a collection bag, where Figure 2A represents a single raw oblique back-illumination capture serving as a representative SCqOBM model input, and Figure 2B corresponds to the matched reconstructed qOBM image using four-captures, which serves as ground truth. Figure 2C and Figure 2D depict the SCqOBM and TCqOBM reconstructions, respectively, which show nearly identical structure as the ground truth. Image insets, in Figure 2E-G, provide a close-up view of an example red blood cell reconstructed using qOBM (ground truth), SCqOBM, and TCqOBM, respectively. Remarkably, all reconstructions show identical qualitative structure. Line segments in Figure 2H-I show that the quantitative refractive index (RI) values are also in excellent agreement.^1^

**Fig. 2.**
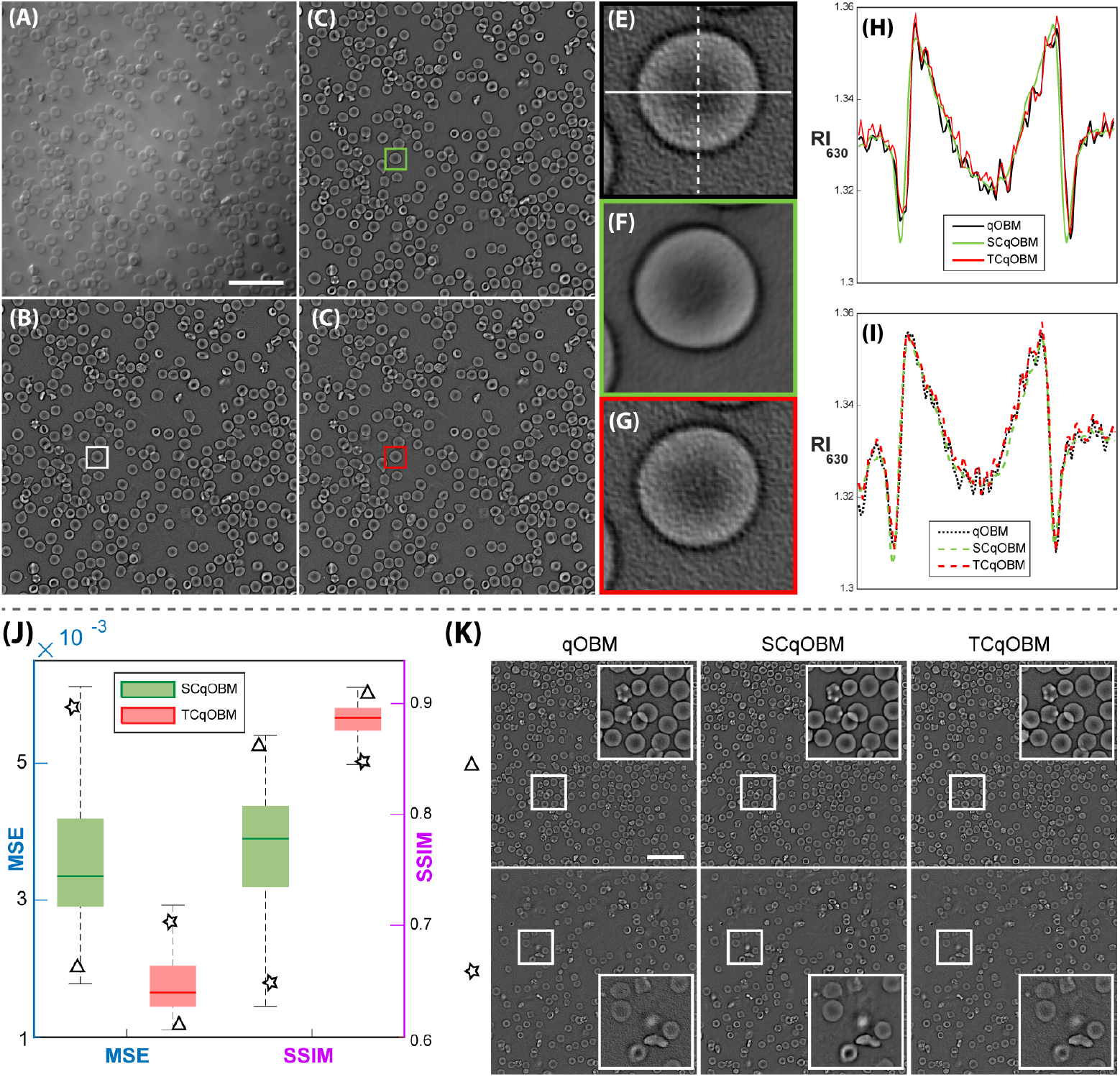
SCqOBM and TCqOBM umbilical cord blood reconstruction results. (A) Representative OBM capture of blood cells taken with 630 nm light. (B-D) qOBM, SCqOBM, and TCqOBM reconstructions of the same image show in (A), respectively. (E-F) qOBM, SCqOBM, and TCqOBM close-ups of regions of interest marked in (B-D), respectively. (H) Horizontal and (I) vertical profiles of the red blood cell from (E-F). The exact regions of the plotted profiles are identified in (E). (J) Box plot showing the MSE and SSIM distributions of the single capture and two capture qOBM modalities. (K) Representative images organized from higher (triangle) to lower (star) performance levels. Scale bar represents 50 *µm*.

To further examine the quantitative reconstruction performance of TCqOBM and SCqOBM, we evaluate the test set mean square error (MSE) and structural similarity index measure (SSIM). Figure 2J-K show the results where the average MSE and SSIM for the TCqOBM reconstruction are 0.0018 and 0.89, respectively, and 0.0036 and 0.77 for SCqOBM. Both of these metrics show the excellent quantitative accuracy of both TCqOBM and SCqOBM for reconstructing these relatively simply structures, with the TCqOBM slightly out performing SCqOBM (more on this below). We also note that SSIM is a measure of structural differences, and it is highly susceptible to disparities in noise, which accounts for the observed decline in SCqOBM SSIM values. Figure 2 K shows three representative reconstructions with varying levels of measured performance, highlighting that reconstructions with well in focus structures show high fidelity quantitative reconstructions, while images with out-of-focus content tend to yield inferior results. Improvements can be attained with more extensive examples of images with out-of-focus content in training.

The observed MSE and SSIM values indicate that TCqOBM achieves higher reconstruction fidelity compared to SCqOBM. This can be potentially attributed to the additional directional information provided by the two captures with orthogonal shear information, which aids in more accurate phase retrieval in all directions in TCqOBM compared to SCqOBM (see Methods Section). However, SCqOBM still performs remarkably well, and has clear advantages in applications requiring rapid data acquisition. For these types of idealized samples, comprising mostly of symmetrically shaped biconcave RBCs, future work could explore further optimization of the deep learning model to enhance the SCqOBM performance.

#### 2.1.2 *Ex-vivo* bulk fresh brain tissue

Next, we investigate the performance of TCqOBM and SCqOBM in more complex samples, comprising a dataset of images of rat brain tissue as described in Ref. [20]. The DL models for this data set had an identical structure and training protocol as that used for the blood models described above, but the models were trained independently. The dataset employed for this process consisted of images acquired with a 60x (0.7 NA) objective, using 720 nm LED illumination for all 4 acquisitions in the original qOBM dataset.

The rat brain dataset presents a drastically more complex challenge compared to the blood dataset, characterized by a highly symmetrical structures. This dataset consists of images of fresh rat brains taken from various regions of healthy brain, as well as images of rat brain tumors, together resulting in highly diverse RI distributions. This variability is evident in Figure 3A-L, which displays representative brain tissue images with three very distinct architectures: bulk tumor, tumor margin, and healthy cortex. The bulk tumor tissue exhibits higher refractive index values while the gray matter in the cortex displays finer structures with lower refractive index [20]. We also show a tumor margin region, which combines large tumor cells with high RI values with the finer structures of the surrounding healthy tissue.

**Fig. 3.**
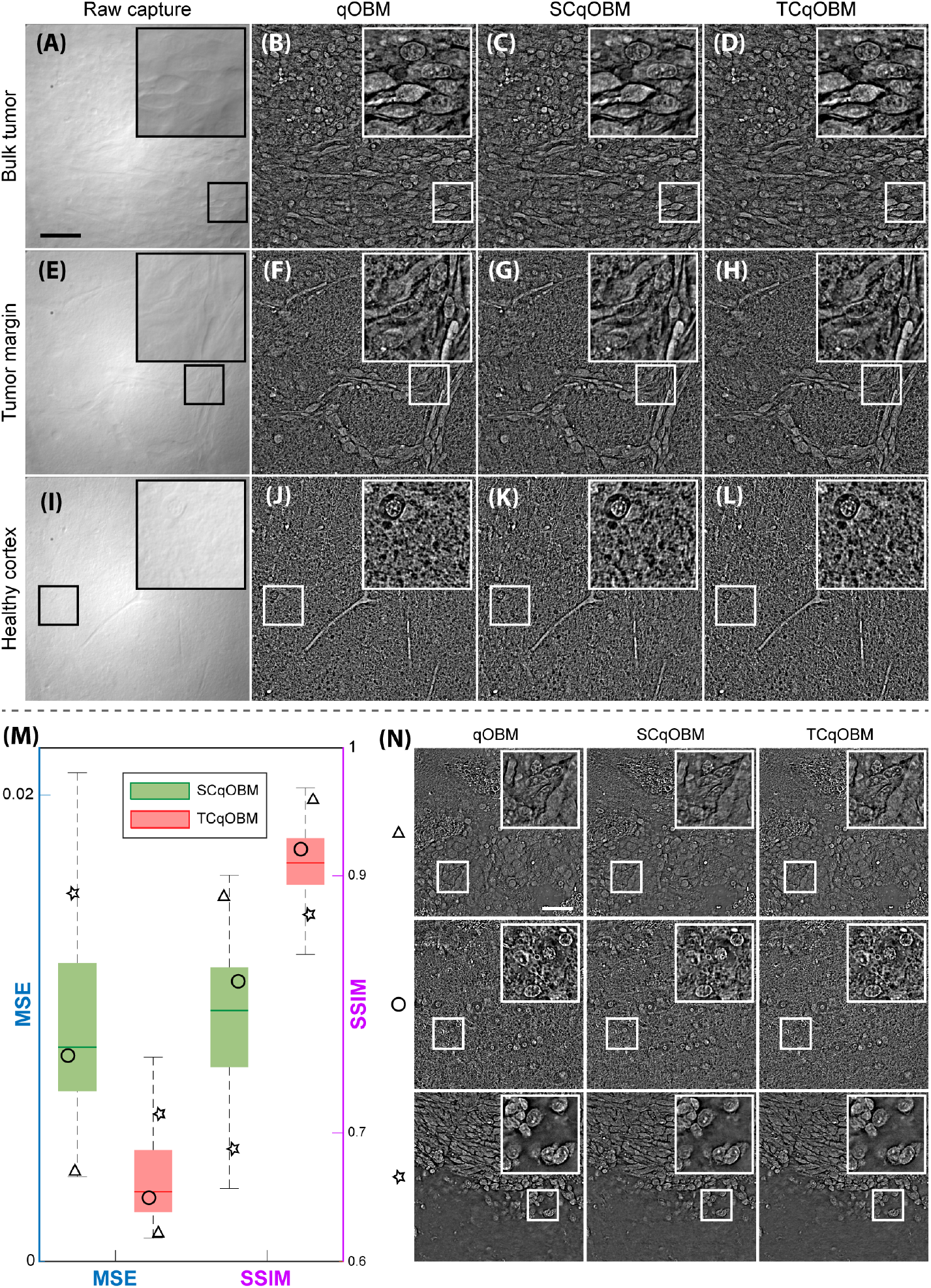
SCqOBM and TCqOBM rat brain tissue. (A-D) Raw capture, qOBM, SCqOBM, and TCqOBM of a tumor region in rat brain. (E-H) Raw capture, qOBM, SCqOBM, and TCqOBM of a tumor margin region in rat brain. (I-L) Raw capture, qOBM, SCqOBM, and TCqOBM of a healthy cortex region in rat brain. (M) Box plot showing the MSE and SSIM distributions of the single capture and two capture qOBM modalities. (N) Representative images organized from higher (triangle) to lower (star) performance levels. Scale bar represents 50 *µm*.

Despite the complexity of the brain tissue, the DL-based phase retrieval for TCqOBM and SCqOBM (Figure 3) excel at recapitulating the quantitative phase information from all types of structures. By examining the performance metrics depicted in Figure 3M, we observe that the SSIM scores for this dataset are better than those for the blood dataset, with an average of 0.8 and 0.91 for SCqOBM and TCqOBM, respectively, while the MSE values are worse for the brain dataset, averaging 0.1 for the SCqOBM and 0.036 for the TCqOBM. This indicates that the tissue structures are recovered remarkably well, but with a slightly higher phase error than the blood images. Nevertheless, these error values remain extremely low and almost negligible in relation to the reconstructed phase values, where the mean absolute percentage error is *<* 0.6% for all the presented frameworks.

Figure 3N illustrates a set of representative rat brain images with varying performance levels. The images identified with a star symbol, indicating worse performance metrics, feature substantial background (tissue-less) regions. It does not come as a surprise that the model’s precision in reconstructing the background is compromised, considering that the training data patches underwent quality assessments to ensure only high-fidelity data with tissue structures were used during training (see Methods Section). Background regions like those shown in the last row of Figure 3N do not contain important information, and thus their accurate reconstruction is not a priority. Importantly, tissue regions, even in images containing significant background, show remarkable agreement with the “ground truth” and are essentially qualitatively indistinguishable from one another. This underscores the huge potential for TCqOBM and in particular SCqOBM to improve imaging speeds while retaining high quantitative fidelity, even when encountering highly heterogeneous tissue samples.

As a final validation of TCqOBM and SCqOBM using static samples, we tested the two models trained on rat brain tissue on images taken of human glioblastoma tissue (discarded from neurosurgery) [30]. While the optical properties of human and rat brain tissues are equivalent, the structures present in these two datasets are quite distinct. As shown in Figure 4, the SCqOBM and TCqOBM models are able to successfully recover the quantitative phase information from human tissues with drastically different structures from those seen in training (rat brain) with remarkable agreement. The MSE and SSIM values of the human tissues are also within the same range as that seen in the test set of the rat brain (Figure 3M). This suggest that the models can perform well even when encountering significantly different structures, which is critically important for broad clinical and other applications.

**Fig. 4.**
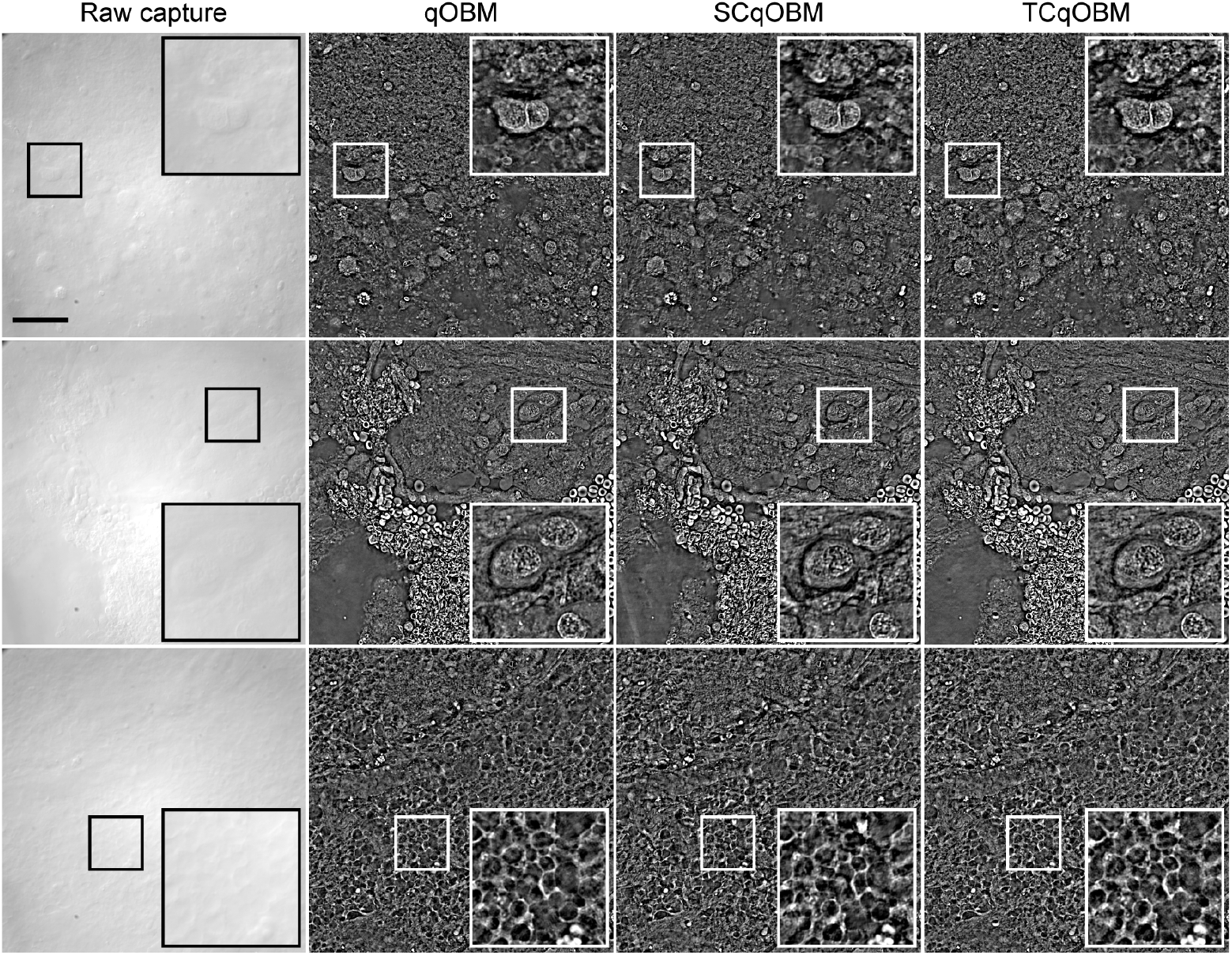
SCqOBM and TCqOBM human brain tissue reconstruction results. (Left to right) a raw capture, qOBM, SCqOBM, and TCqOBM reconstructions of different regions acquired from fresh human glioblastoma specimens discarded from neurosurgery. Scale bar represents 50 *µm*.

#### 2.1.3 Fourier Analysis of TCqOBM and SCqOBM

Before moving to dynamic samples, it is important to gain a better understanding of the differences between qOBM and the two DL-based reconstruction methods, TCqOBM and SCqOBM. For this task, it is most useful to analyze the data in the Fourier domain. Because qOBM uses two sets of orthogonal DPC images to ensue quantitative phase recovery with high fidelity along all directions, the coverages in the frequency domain is isotropic in the transverse direction [12, 15]. For TCqOBM, two captures (one from each set of the two orthogonal DPC images) are used to also ensure information is recovered along all directions, while SCqOBM foregoes acquisition along one direction to enable faster imaging. Figure 5 shows the Fourier domain of raw, qOBM, TCqOBM, and SCqOBM images of blood and brain specimens. Note that a single raw oblique illumination capture has missing information along a narrow region orthogonal to the direction of oblique illumination. As expected, the TCqOBM approach, like qOBM, recovers the full angular information of the phase, thanks to the perpendicular illumination direction of the two captures used in this process. The SCqOBM, however, struggles to recover information not contained in the input image (single raw oblique illumination), but this depends on the sample type. When reconstructing blood cell images, for example the SCqOBM model learns to recover the angular information missing from the raw capture. This is likely due to the highly predictable circular structure of the blood cells. On the other hand, for brain tissue, with far more complex and heterogeneous architecture, SCqOBM is not able to predict/infer the contents of the missing information. Thus, as the lower row of Figure 5 shows, the missing information of the raw single capture is not recovered (nor inferred) in SCqOBM. Interestingly, however, SCqOBM does not attempt to fill in or “hallucinate” structures along those regions; instead, structures along the narrow missing frequency band remain absent.

**Fig. 5.**
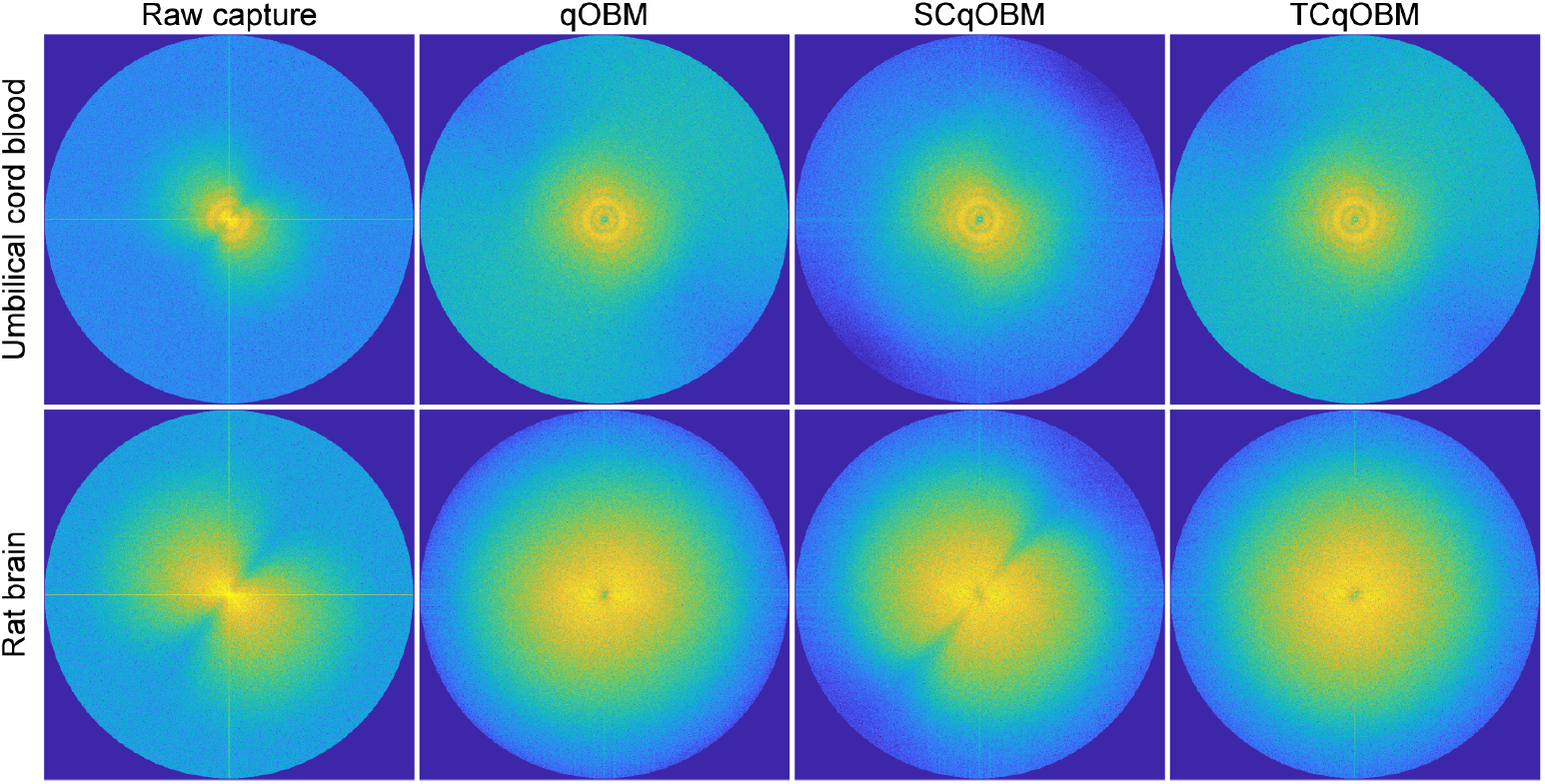
Recovered frequency space through different approaches. Logarithmic representations of the Fourier domain of (left to right) a raw capture, qOBM, SCqOBM, and TCqOBM taken from umbilical cord blood (top) and rat brain tissue (bottom).

### 2.2 Fast qOBM of dynamic *in-vivo* samples

#### 2.2.1 *In-vivo* blood flow analysis in mouse brain

Next we analyze blood flow in a mouse brain using a window chamber model. Here we used a fast frame-rate camera that enabled imaging rates of ∼ 2 kHz to quantify blood flow through blood vessels of varying sizes and all data were processed using the same SCqOBM model trained with images of rat brain tissue. As illustrated in Figure 6A, blood flow rates were quantified using temporal refractive index fluctuations from blood cells flowing along a line path along a vessel. By plotting the refractive index along this line as a function of time, blood cells flowing in the vessel appear as slanted lines, from which speed can be readily calculated. This method for extracting blood flow is similar to M-mode ultrasound imaging [33]. As there will be variations in speed over time for any given particle flowing along the lines’ path, we calculate the slope for all lines by rotating the M-mode image over an angular range and integrating the pixel intensity over distance. The angle at which a line’s summed pixel intensity is lowest represents the slope and thus speed of a particle over that distance. The range of speeds acquired using this method for three vessel sizes (7 *µm*, 20 *µm*, and 80 *µm* in diameter) can be seen in Figure 6C.

**Fig. 6.**
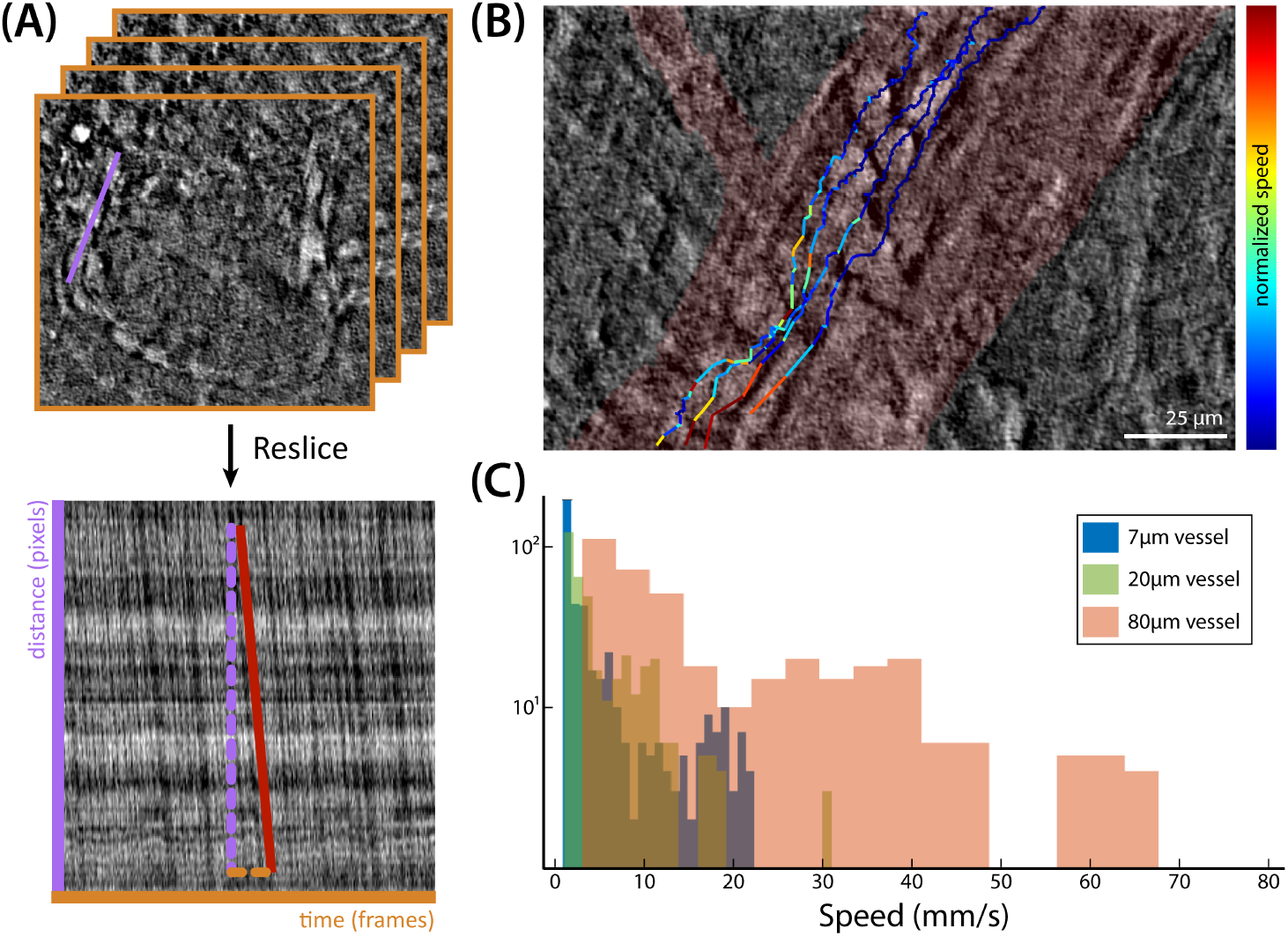
Analysis of blood flow patterns in mouse cranial vasculature. (A) shows the process of extracting the flow speed within a vessel. A line along the direction of flow is drawn and plotted against time to create an image. The vessel’s flow speeds can be calculated from the slope of the lines in the distance vs time plot. (C) demonstrates a histogram of the calculated speeds from these slope lines shows flow speeds across 7*µm*, 20*µm*, and 80*µm* vessels. (B) shows the path of four white blood cells across a 40*µm* vessel is shown. The trajectories are encoded by the normalized speed of the WBC along its path.

Blood flow rates ranged from 1.1 mm/s in the smallest vessels (7*µm*) to 66.4 mm/s in the center of the largest vessels (80*µm*). Moreover, we observed physiologically expected changes in speed across the width of a vessel [34], where the speeds at the edge of an 80 *µm* vessel are less than the speeds at its center forming a characteristic Poiseuille flow profile, as seen in Figure A2B. Additionally, differences in flow rates between similarly sized feeder vessels were observed. Vessels of the same size (16*µm*) that fed into a slightly larger vessel (32*µm*) exhibited higher average speeds at 10.0 mm/s, compared to a vessel that fed into a significantly larger vessel (80*µm*) with an average speed of 6.7 mm/s, shown in Figure A2D.

Finally, slower moving leukocyte rolling across larger vessels (*>* 40*µm*) were also observed and characterized. For example, the path and speed of four rolling leukocyte can be seen in Figure 6B; live speed tracking for each of the leukocytes can be seen in Supplementary Videos (Video 1-4). These results have important implications, as leukocyte adherence to vascular endothelium, rolling, and eventual extravasation into the surrounding tissues marks an important immune response[35].

In fact, as the mouse approached end of life throughout the imaging session, the density of adherent leukocytes increased progressively. The difference in leukocyte adherence between an earlier and later time point can be compared between Supplementary Video 5 and Video 6, respectively.

### 2.3 Fast volumetric imaging of human skin in vivo

As a final demonstration of the unique capabilities of SCqOBM, we demonstrate near video rate volumetric refractive index tomography of human skin (arm) in vivo. In order to perform 3D imaging at a high speed, we added a z-axis piezo-stage in our system (see Methods section). Raw intensity stacks are first converted to 2D phase sections with SCqOBM, and then the 3D image stack is processed using a separate DL-based 3D algorithm that improves the refractive-index quantitative fidelity and SNR (see Ref. [36] for details). The resulting volume provide clear cellular and sub-cellular contrast. From the surface down to ∼ 100*µm*, one can observe the stratum corneum, stratum granulosum, stratum spinosum, and stratum basale (Figure 7A). At greater depths, between ∼ 100 *−* 150*µm*, blood cells moving through capillaries in the dermis can be observed (Figure 7A-B).

**Fig. 7.**
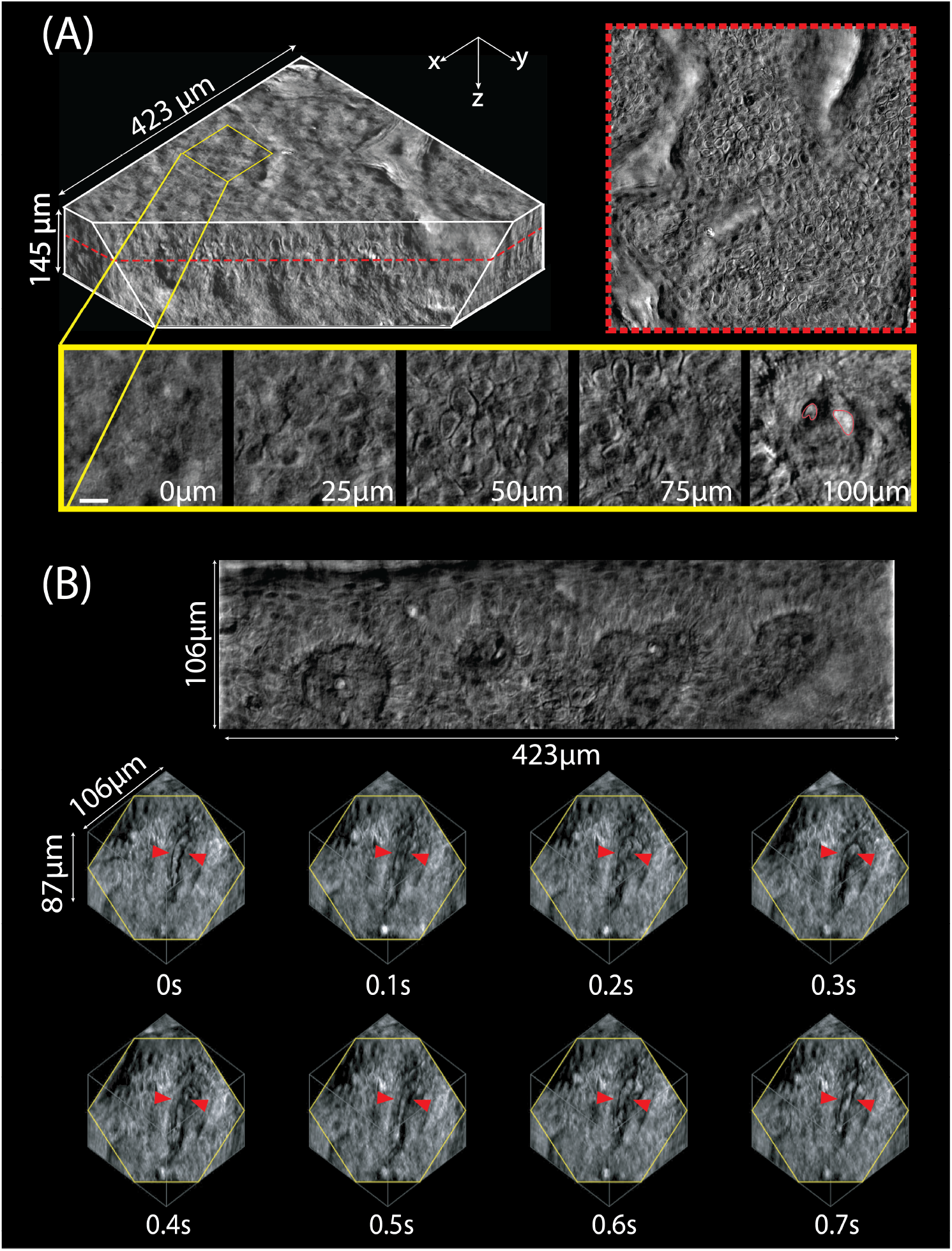
(A) Rendered volume spanning 423 µm × 423 µm × 145 µm (x-y-z) acquired with 1.45 µm axial steps (100 sections). The red dashed line highlights the basal-layer region whose full field of view is displayed on the right. Five magnified insets (yellow squares, bottom) reveal successive epidermal and dermal layers—from the stratum corneum down to the papillary dermis. Two individual red blood cells are marked in red. (B) High-speed acquisition of a reduced field of view (512 × 2048 pixel) captured at 10 VPS. Consecutive reconstructions spaced by 100 ms illustrate RBC flow through different planes.

Imaging parameters can be flexibly tuned based on the needs of a given application. Specifically, the active sensor area and number of z-planes can be adjusted depending on the desired image conditions. Importantly, in our system the total number of voxels recorded per second - here 1.25 Giga-voxels per second - remains relatively constant allowing users to tune the spatial extent and temporal resolution without sacrificing overall information throughput. For example, Figure 7A shows a volume of 2048 × 2048 × 100 voxels, corresponding to 423 *µm* × 423 *µm* × 145 *µm*, acquired using a 40 ×, 0.6 NA objective. In this mode, we can capture relatively larger volume at a modest rate of three volumes per second (VPS), shown in 3D in Supplementary Video 7. Red blood cells can be visualized at this speed in Supplementary Video 8. In Figure 7B, on the other hand, we restrict the field of view to one quarter of its original area (512 × 2048 pixels) and sample 40 axial planes, allowing acquisition at 10 VPS. In this faster imaging mode, we can visualize blood flow in the human arm in 3D as the cells loop through the capillaries at the top of the dermis (red arrows). The figure captures 4 capillaries in one plane (top row) and a 3D volume acquired from one of them (cropped for visualization) at different times (middle and bottom rows). Demonstration of the red blood cells flowing at 10 VPS can be found in Supplementary Videos 9-11. With an effective throughput of 1.25 Gpixels per second, SCqOBM achieves acquisition speeds comparable to state-of-the-art rapid volumetric light-sheet microscopes.

## 3 Methods

### 3.1 qOBM system configuration

The configuration of the traditional qOBM microscope is shown in Figure 1A. Its geometry is that of a conventional inverted microscope with a modified illumination scheme. Samples are sequentially illuminated in epi-mode with four LED light sources coupled into multimode fiber optics (1 mm in diameter). The LED wavelength varied for different applications (527 nm, 650 nm, 720 nm, or 850 nm). The fibers are positioned around the objective lens 90-degrees from one another and 45 degrees from the optical axis on a custom 3D printed adapter. When light from an LED source is deployed through one of the fibers into the thick sample, it undergoes multiple scattering, causing some of the photons to change trajectory and effectively producing a virtual light source within the thick object, emulating a transmission source with a slight offset to the optical axis.

In qOBM, lateral variations in the refractive index at the focal plane cause light to refract towards or away from the objective’s acceptance angles (defined by its numerical aperture), resulting in phase contrast. By subtracting images captured from diametrically opposed fibers, differential phase contrast (DPC) is generated, which helps to eliminate out-of-focus content, enabling tomographic sectioning (Figure 1F,G,I). To produce quantitative phase contrast, two orthogonal DPC images (derived from four raw captures) are processed. This involves modeling the angular distribution of multiple-scattered light within the sample (mcxlab run on MATLAB), which acts as an effective, transmission-like illumination source. The transfer function can also be measured experimentally [36]. The resultant optical transfer function estimate (or measured) is then used to extract quantitative phase information via Tikhonov regularized deconvolution (Figure 1H,I) [12, 15, 17].

SCqOBM and TCqOBM image acquisition is carried out with a similar setup to traditional qOBM, but with fewer illumination fibers — one or two, respectively. In TCqOBM, the illumination fibers are positioned at a 90° angle from each other to provide perpendicular illumination between the two raw captures obtained. This positioning ensures that the frequency domain contains relevant information from all angles.

The image reconstruction for SCqOBM and TCqOBM is performed using a deep neural network, with the raw acquired data as the input and the quantitative phase image as the output. Detailed information on the model architecture and training process is provided in the following sections.

### 3.2 Generative adversarial networks for qOBM image reconstruction

Operations involved in the translation of images from one domain to another can be considered as a segmentation task with a very larger number of potential class labels. U-Nets can be a suitable candidates to address these types of problems, but, when trained traditionally, they often overlook the target image distribution, resulting in the generation of images that lack natural visual characteristics. To mitigate this issue, the inclusion of a referee mechanism during training can be employed to assess whether the generated images adjust to an expected distribution of “natural images”. This can be achieved through the adoption of a U-Net-based generative adversarial network (GAN) framework (often referred to as *Pix2Pix*), where the U-Net functions as the generator responsible for image translation, while a classifier DL model, known as the discriminator, determines whether the generator’s outputs fall within the expected distribution of real images [37, 38]. The training SCqOBM and TCqOBM models follow this approach.

Within a GAN framework, the primary objective is to determine a generator transformation function, *G*_*X*_ : *X → Y*, to effectively map the input dataset *X* to the output dataset *Y*. The success of the generator depends on the distribution of *G*_*X*_ (*X*) being indistinguishable from samples drawn from the distribution *Y*. During the training process, the discriminator becomes increasingly better at distinguishing between real and generated images, while the generator improves its translated image generation. The ideal outcome of the process is for the generator to generate an output that the discriminator can not flag as fake. This objective can be summed up in the loss function of the GAN, which is given by

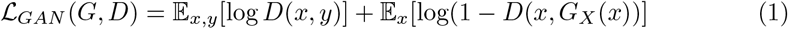

A common approach to this framework includes mixing a distance loss (such as L1 or L2) to improve the generator’s output. As such, the final loss for the SCqOBM framework yields

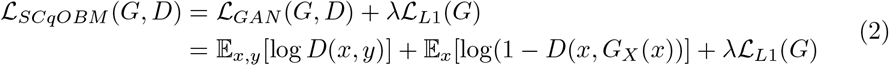

where *ℒ*_*L*1_(*G*) is the L1 distance loss between the ground truth and generated output, and *λ* is a weight factor between 0 and 100 to scale the importance of the distance loss over the GAN loss. It is worth noting that conditional U-Net GAN like the one described here, require pixel level matching between the input data and the ground truth. This requirement was satisfied for the training of all models presented in this chapter.

In order to train the models, the datasets were divided to independent training and testing sets. For computational simplicity, each image was cropped into sections of 256 px × 256 px (∼ 35 × 35 *µm*). The training was performed with batches of 8 patches each. In each training iteration, the model weights were updated using the data from a single batch. For each model, ∼ 120,000 image sections were used for training, and ∼ 20,000 independent ones for testing. All models were solely trained with data taken with a 60x objective (0.7 NA).

The GAN training process is illustrated in Figure A1, where the ground truth *y* is a qOBM image reconstructed with four OBM captures through Tikhonov deconvolution (see Figure 1I). Meanwhile, the input consists of one or two OBM captures, as described below. The raw images are fed through the generator, which attempts to produce a map that matches the target (i.e., the ground truth image). The discriminator then attempts to guess whether the generated images are real or fake. Consequently, the discriminator weights are updated through back propagation based on its accuracy in the labeling of real (*y*) and fake (*G*_*X*_ (*x*)) data. Next, the generator is updated based on two primary criteria: (1) its ability to deceive the discriminator and (2) its capability to produce an image that closely resembles the ground truth. All model training, validation, and processing was run using the PyTorch framework. The models were trained using an NVIDIA RTX 3090 GPU, with the Adam optimizer (learning rate = 0.002, beta1 = 0.5) for 20 epochs. Once trained, the model takes 3.5 ms to process the input images to TC-/SC-qOBM (2048 px X 2048 px image, using an Nvidia GeForce 3090 Ti GPU).

#### 3.2.1 Two capture qOBM

To evaluate the ability of GAN to retrieve the quantitative phase from OBM images, we first studied the DL-based reconstruction from two raw captures by training a GAN, which we name TCqOBM. The two captures are taken from illumination sources positioned 90 degrees from one another, and thus they contain contrast in perpendicular directions. Therefore, the information regarding RI variations in all directions is contained within those two captures. The captures were concatenated in the third dimension, such that the TCqOBM model produces a single channel (grayscale) output from a two channel input, where each channel represents a distinct angle of illumination.

#### 3.2.2 Single capture qOBM

The SCqOBM follows a comparable approach, but with a single OBM image as the input. Throughout this process, the offset angle between the illumination shear direction and the detector remained consistent, that is, the raw captures used to train the SCqOBM models were obtained by positioning the illumination fiber at roughly the same position relative to the camera. This approach ensures that the relative direction of the illumination gradient in the input image remains approximately constant, and the model can more effectively interpret the data. (Because the shear angel is relative, this can also be achieved by digitally rotating the image.)

We assess the performance of these DL-based phase retrieval algorithms by conducting experiments involving the reconstruction of two biomedical specimens: umbilical cord blood [14] and fresh, thick rat brain tissue [20]. The results are presented in the following sections.

### 3.3 Sample preparation and imaging conditions

#### 3.3.1 qOBM imaging of umbilical cord blood and freshly excised tissue

The umbilical cord blood units (CBUs) were obtained from the Carolinas Cord Blood Bank at the Duke University Medical Center and imaged in the Optical Imaging and Spectroscopy (OIS) Lab at the Georgia Institute of Technology. Only research CBUs which did not meet storage criteria due to low collection volumes were used. Samples were imaged within 48 hours of collection. More information about the samples and imaging process can be found in [14].

All animal experimental protocols were approved by Institutional Animal Care and Use Committee (IACUC) of the Georgia Institute of Technology and Emory University. Mouse and rat brains were imaged withing 24 hours of collection. More information about the samples and imaging process can be found in [20].

#### 3.3.2 *In-vivo* mouse brain imaging

All experimental procedures were approved by the Institutional Animal Care and Use Committee (IACUC) at the Georgia Institute of Technology. 3-5 month old C57Bl6/J mice (1 male, 2 female) were anesthetized under isoflurane (3% induction, 1.5-2% maintenance). Following induction of anesthesia, mice were placed in a stereotaxic apparatus to stabilize the head and the body temperature was maintained at 37° C with a heating pad and rectal temperature probe. Ketoprofen (5 mg/kg) and atropine (0.05/mg/kg) were administered subcutaneously. The eyes were protected with ophthalmic ointment and 2% lidocaine was injected under the scalp. An incision of the skin was made to expose the cranium and the fascia was removed by gently scraping the skull surface. A stainless steel or aluminum headplate was adhered to the skull with cyanoacrylate glue (Vetbond) and dental cement (C&B Metabond). A 1-2 mm diameter craniotomy was performed over the left somatomotor cortex approximately 1 mm posterior and lateral of Bregma by thinning the bone around the craniotomy site with a dental drill and removing the central bone flap with fine forceps under a drop of artificial cerebrospinal fluid (ACSF, in mM: 150 NaCl, 4 KCl, 10 HEPES, 2 CaCl2, 1 MgCl2, pH 7.4, 305-310 mOsm). The dura was left intact. A 5 mM circular glass coverslip was placed over the craniotomy such that it remained submerged in ACSF and the edges of the coverslip were sealed with silicone polymer (Kwik-cast). The mouse was transferred to the imaging rig and head-fixed by clamping the headplate. Isoflurane anesthesia (1-2%) and temperature were maintained during the transfer and imaging session with a nosecone and rectal temperature-controlled heating pad. Respiratory rate and toe-pinch response was monitored throughout to ensure a surgical plane of anesthesia. Upon completion of imaging experiments, the mouse was deeply anesthetized under isoflurane (5%) and euthanized by cervical dislocation.

#### 3.3.3 *In-vivo* human arm imaging

To enable in-vivo human arm imaging (using healthy volunteers), we use two different system orientations to achieve maximal stability at different placements of the arm. For posterior arm imaging, we utilize the upright system orientation such that the objective points downwards. A glass panel is placed between the objective and the arm, and stability is achieved either by slight compression between the glass panel and a flat surface or by an external arm rest. For anterior arm imaging, we invert the system orientation such that the objective points upward. The arm is then imaged through a glass slide, and stability is achieved by slight external compression of the imaged arm. To enable deeper penetration into the skin, a single LED with a wavelength of 840nm is used.

In order to perform 3D imaging at a high speed, we added a z-axis piezo-stage to our system. The queenstage NanoScan OP400 has a 400*µm* closed loop travel range, allows us to easily and rapidly change the imaging plane of our objective and image from the surface to inside of the tissue within milliseconds. The step size is calculated so that the 3D model [36] trained for volume can have the best reconstruction performance. For data processing, SCqOBM is applied first to transform the raw capture to phase images, then, we input a z-stack of phase images into the 3D model, which provides a higher fidelity refractive index tomographic reconstruction.

## 4 Discussion

In this work we have introduced SC- and TC-qOBM, as a means to significantly improve the imaging speed of qOBM. We demonstrated the TC-qOBM recovers phase information that is nearly identical (qualitatively and quantitatively) to traditional qOBM (processed using four captures and a deconvolution). SCqOBM, on the other hand, maximizing the imaging speed of qOBM and produces qualitatively identical results to qOBM, but quantitatively lacks structural information along a narrow range of spatial frequencies that lie orthogonal to the oblique illumination light source. Nevertheless, outside this narrow range of spatial frequencies, relatively high quantitative fidelity was still maintained in SCqOBM, even in complex samples such as the brain.

The DL-based phase retrieval frameworks were first validated using static samples where ground truth (traditional qOBM images) was known *apriori* without movement artifacts. Importantly, we showed that the models trained using mouse brain tissue were readily generalizable to other tissue types, including human brain and skin. Though, improvements can be expected with a tissue-specific model. Then, to demonstrate the power of this imaging pipeline, we imaged and quantitatively analyzed blood flow mouse brains in-vivo. Finally, we also demonstrated fast volumetric skin imaging in humans, where clear cellular and subcellular detail can be observed. Fast volumetric imaging acquired up to 1.25 Gigavoxels per second enabled 3D visualization of blood flow through capillaries in the dermis of healthy volunteers.

Compared to traditional qOBM, SCqOBM (and TCqOBM) do have some tradeoffs. First, high-speed cameras with low dark noise and wide dynamic range are important to achieve the desired imaging rates and maintain high signal-to-noise ratio (SNR). Such cameras are bulkier and more expensive. Also, because of the higher frame-rates and shortened integration times, higher power LEDs (¡150 mW at the sample) are necessary to ensure sufficient light is collected. However, tissue damage is still not a concern given use of near infrared light (850 nm) with diffused illumination. SCqOBM represents a significant advancement in quantitative phase imaging and label-free imaging, offering a fast, simple, tomographic, and accessible means of recovering high contrast, cellular and subcellular information from thick, scattering samples. The demonstrated framework enables imaging speeds comparable to state-of-the-art light sheet microscopy, with penetration depths comparable to label-free nonlinear microscopy, while offering the same rich structural information provided by QPI. Its potential applications in biomedical research and medicine are vast, ranging from non-invasive blood flow analysis to real-time *in-vivo* volumetric imaging and more. We expect SCqOBM to a valuable tool for advancing our understanding of complex biological systems and improving patient care.

## 5 Supplementary information

Video 1: Speed-encoded leukocyte 1 tract shown in Fig. 6B as it flows through a mouse blood vessel. Speed is encoded in mm/s, and video is played back at 20 fps (originally captured at 500 fps).

Video 2: Speed-encoded leukocyte 2 tract shown in Fig. 6B as it flows through a mouse blood vessel. Speed is encoded in mm/s, and video is played back at 20fps (originally captured at 500 fps).

Video 3: Speed-encoded leukocyte 3 tract shown in Fig. 6B as it flows through a mouse blood vessel. Speed is encoded in mm/s, and video is played back at 20 fps (originally captured at 500 fps).

Video 4: Speed-encoded leukocyte 4 tract shown in Fig. 6B as it flows through a mouse blood vessel. Speed is encoded in mm/s, and video is played back at 20 fps (originally captured at 500 fps).

Video 5: Blood flow through mouse vessel with few leukocytes visible captured at 500 fps, shown at 100 fps.

Video 6: Blood flow through mouse vessel with many more leukocytes visible later in acquisition captured at 500 fps, shown at 100 fps.

Video 7: Human skin volume shown in Figure 7A rotated and sliced in 3D.

Video 8: Axial pan through skin volume shown in 7A with highlighted red blood cells shown.

Video 9: 10 vps (played at 10 fps, repeated 10 times) skin blood flow data shown as the 87 um sectioned volume in Fig. 7B, with blood flowing through different Z-planes

Video 10: 10 vps (played at 10 fps, repeated 10 times) skin blood flow data (Fig. 7B) shown as a plane in more superficial parts of the epidermis

Video 11: 10 vps (played at 10 fps, repeated 10 times) skin blood flow data (Fig. 7B) shown as a plane in deeper parts of the epidermis (20 um deeper)

## Declarations

## 5.1 Acknowledgments

We acknowledge the following funding sources:

Burroughs Wellcome Fund: CASI BWF 1014540 (F.E.R.)

National Institute of General Medical Sciences: R35GM147437 (F.E.R.)

Georgia Research Alliance (F.E.R.) Georgia Institute of Technology: (F.E.R.)

## 5.3 Author Contributions

P.C.C. and F.E.R. conceived the research, P.C.C. imaged the blood data and ex-vivo rat brain data, P.C.C. and S.R. imaged the in-vivo brain data, A.L, M.LE and B.H. prepared the animals for in-vivo brain data collection, S.R. and Z.L. imaged and processed the in-vivo human skin data P.C.C. and N.K. developed the deep learning pipeline and trained the models, P.C.C., S.R., and Z.L. performed the result analysis and statistical study. P.C.C., S.R., Z.L., and F.E.R. prepared the manuscript, and all authors contributed to the manuscript. F.E.R. supervised the research.

## 5.4 Data and Code availability

All the data and methods that support this work are present in the main text and the Supplementary Information. The deep learning models in this work employ standard libraries and scripts that are publicly available in PyTorch.

## 5.5 Ethics Approval

All experimental procedures were approved by the Institutional Animal Care and Use Committee (IACUC) at the Georgia Institute of Technology (Protocol A100171).

## Appendix A Extended Data

**Fig. A1.**
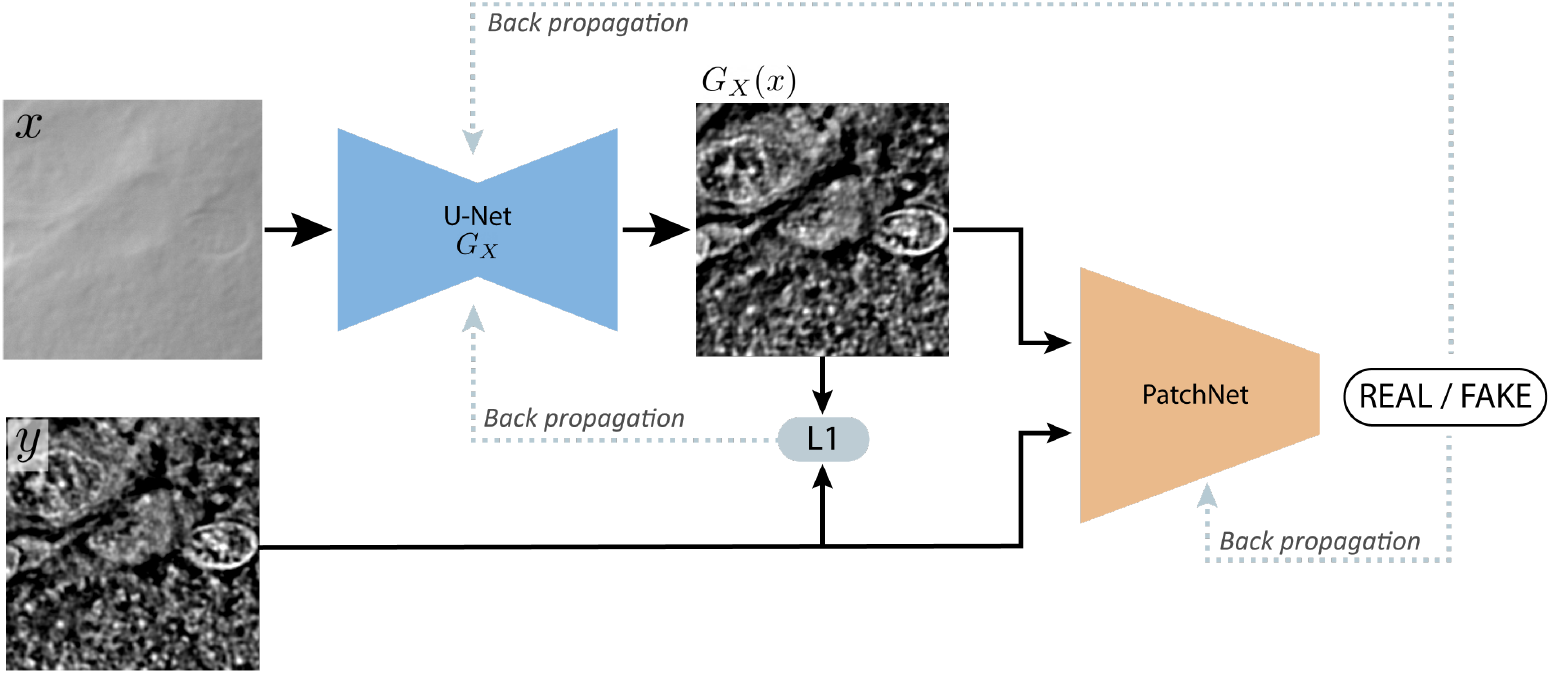
SCqOBM GAN architecture and training process. A raw input image (*x*) of ∼35 *µm × ∼* 35 *µm* is fed through the U-Net generator, which reconstructs a prediction of the quantitative phase (*G*_*X*_ (*x*)). The prediction, along with the ground truth image (*y*) are then passed through the PatchNet, which serves as discriminator. The discriminator loss is computed from its prediction and its weights are updated through back-propagation. The weights of the generator are updated based on the combination of the L1 distance loss between the prediction and the ground truth and the outputs of the discriminator.

**Fig. A2.**
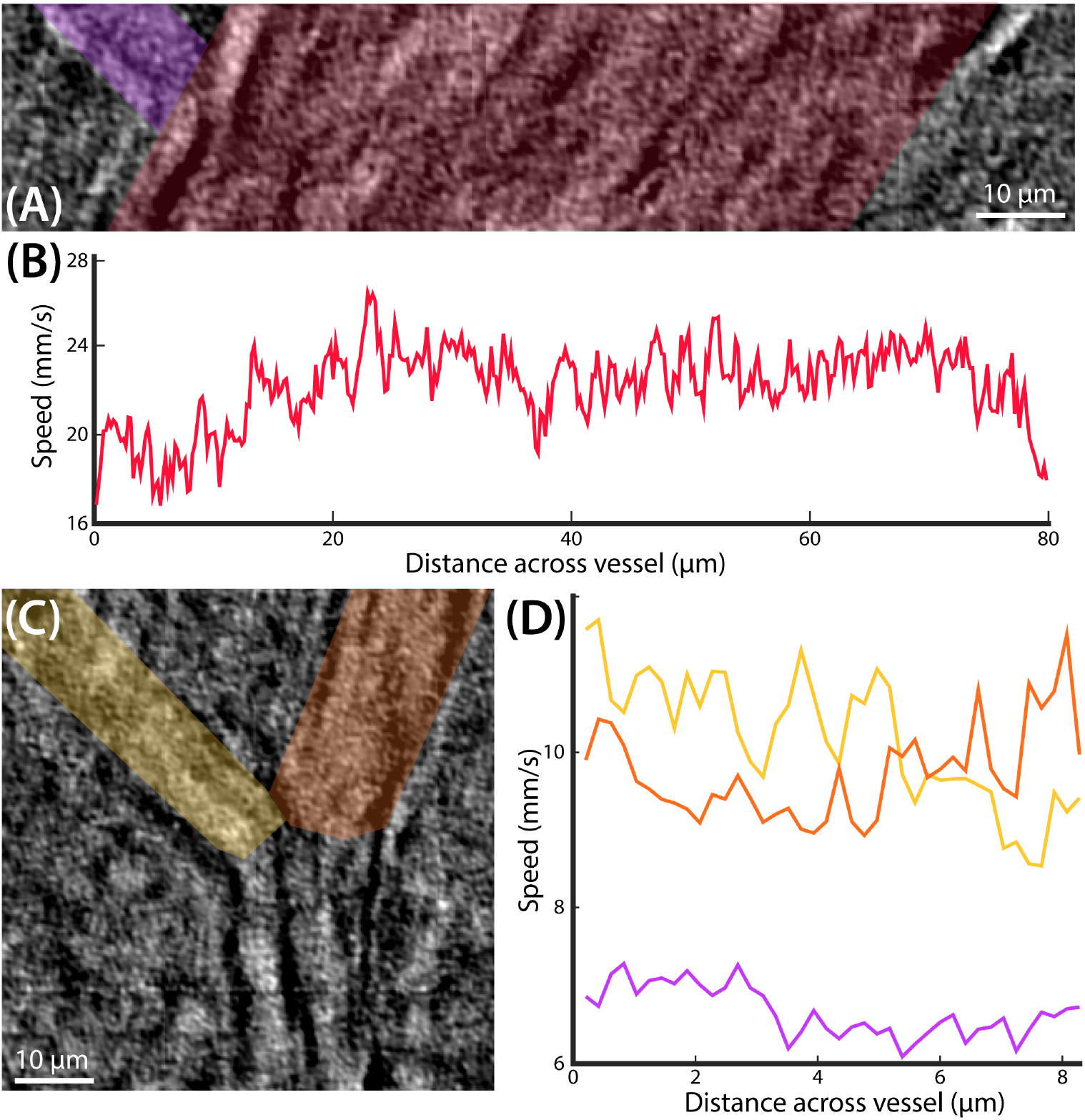
Blood flow speeds across mouse cranial vasculature. (A) shows an 80*µm* vessel (highlighted in magenta) and its 12*µm* feeder vessel (highlighted in purple). (B) highlights the profile of the blood flow across the 80*µm* vessel, which demonstrates a Poiseuille-like profile. (C) shows a 32*µm* vessel with a left 16*µm* feeder vessel (highlighted in yellow) and a right 16*µm* feeder vessel (highlighted in orange) (D) demonstrates the flow profile across the three feeder vessels shown in (A) and (C).

## Footnotes

1 Note that both the ground truth (4-capture) qOBM image and the TCqOBM image show lower SNR than the single capture SCqOBM image - this is a result of the use of the green light in the former two reconstructions, which results in lower SNR due to significant hemoglobin absorption. The SCqOBM, which only uses the higher SNR red capture, results in a higher SNR reconstruction.

