## Supplementary videos for "Single Capture Quantitative Oblique Back-Illumination Microscopy"

Video 1: Speed-encoded leukocyte 1 tract shown in Fig. 6B as it flows through a mouse blood vessel. Speed is encoded in mm/s, and video is played back at 20fps (originally captured at 500fps).
Video 2: Speed-encoded leukocyte 2 tract shown in Fig. 6B as it flows through a mouse blood vessel. Speed is encoded in mm/s, and video is played back at 20fps (originally captured at 500fps).
Video 3: Speed-encoded leukocyte 3 tract shown in Fig. 6B as it flows through a mouse blood vessel. Speed is encoded in mm/s, and video is played back at 20fps (originally captured at 500fps).
Video 4: Speed-encoded leukocyte 4 tract shown in Fig. 6B as it flows through a mouse blood vessel. Speed is encoded in mm/s, and video is played back at 20fps (originally captured at 500fps).
Video 5: Blood flow through mouse vessel with few leukocytes visible captured at 500fps, shown at 100fps. 
Video 6: Blood flow through mouse vessel with many more leukocytes visible later in acquisition captured at 500fps, shown at 100fps.
Video 7: Human skin volume shown in Figure 7A rotated and sliced in 3D.
Video 8: Axial pan through skin volume shown in 7A with highlighted red blood cells shown. 
Video 9: 10vps (played at 10fps, repeated 10x) skin blood flow data shown as the 87 um sectioned volume in Fig. 7B, with blood flowing through different Z-planes
Video 10: 10vps (played at 10fps, repeated 10x) skin blood flow data (Fig. 7B) shown as a plane in more superficial parts of the epidermis
Video 11: 10vps (played at 10fps, repeated 10x) skin blood flow data (Fig. 7B) shown as a plane in deeper parts of the epidermis (20 um deeper)
